# Community-level facilitation by macroalgal foundation species peaks at an intermediate level of environmental stress

**DOI:** 10.1101/091777

**Authors:** Ricardo A. Scrosati

## Abstract

This paper examines how community-level facilitation by macroalgal foundation species changes with environmental stress. In rocky intertidal habitats, abiotic stress (mainly due to desiccation and thermal extremes during low tides) increases sharply with elevation because of tide dynamics. A previous study done on the Atlantic coast of Nova Scotia (Canada) showed that, at low elevations, where conditions are benign because low tides are brief, fucoid algal canopies (*Ascophyllum nodosum* and *Fucus* spp.) do not affect the structure of benthic communities. However, at middle and high elevations, where low tides last longer, fucoid canopies limit abiotic extremes near the substrate and, in that way, increase the richness of benthic communities. Richness was measured as the number of benthic algal (except fucoids) and invertebrate species found in replicate quadrats. Using the published data from that study, the present study compares the intensity of facilitation and its importance (relative to all other sources of variation in richness) between middle and high elevations, which represent intermediate and high stress, respectively. Facilitation intensity was calculated as the percent increase in benthic richness between quadrats with low canopy cover and quadrats with high canopy cover, while the importance of facilitation was calculated as the percentage of observed variation in richness that was explained by canopy cover. Data for a total of 688 quadrats surveyed along 350 km of coastline were used. The analyses revealed that both the intensity and importance of facilitation were greater at middle elevations than at high elevations. As canopies were previously found not to affect benthic communities at low elevations, this study indicates that the facilitation–stress relationship viewed at the community level is unimodal for this marine system. Such a trend was already found for some terrestrial systems involving canopy-forming plants as foundation species. Thus, this unimodal pattern may be ubiquitous in nature and, as further studies refine it, might help to predict community-level facilitation depending on environmental stress.

## Introduction

In ecology, facilitation refers to the improvement of species performance caused directly or indirectly by another species (Bruno et al. 2003, Bulleri et al. 2016, Michalet & Pugnaire 2016). Common facilitators are organisms that ameliorate abiotic conditions in environmentally stressful habitats. Examples are alpine cushion plants, which protect smaller plants from cold and wind (Ballantyne & Pickering 2015), desert shrubs, which locally decrease heat and water loss (Pugnaire et al. 2011, Ruttan et al. 2016), and intertidal algae, which limit benthic thermal stress and desiccation at low tide (Bertness et al. 1999, Beermann et al. 2013). The possession of extensive canopies is central to the ability of such species to positively affect others. Because of their influence on entire communities through those mechanisms, those organisms are often referred to as foundation species (Altieri & van de Koppel 2014).

The intensity of facilitation by canopy-forming species depends on the degree of environmental stress. Studies in aquatic and terrestrial communities have consistently found that positive effects are common under abiotically stressful conditions but weak or absent under benign conditions (He et al. 2013). That is a central component of the stress gradient hypothesis (Bertness & Callaway 1994), which has frequently been used to assume a continuous increase in facilitation intensity with abiotic stress (Maestre et al. 2009). More recent studies have began to evaluate if the intensity of facilitation may actually often have a unimodal relationship with stress, especially if facilitation is evaluated at the community level (Michalet et al. 2006, Brooker et al. 2008, Holmgren & Scheffer 2010). An important reason for such a pattern could be that, under extreme stress, facilitators may be unable to ameliorate conditions strongly enough for many species (Holmgren & Scheffer 2010, He & Bertness 2014, Michalet et al. 2014). This paper investigates the facilitation–stress relationship at the community level using data from rocky intertidal systems.

The intertidal zone is the area of the coast between the highest and lowest tidal levels. As low tides become longer with intertidal elevation, biological desiccation and thermal extremes increase with elevation because of the longer exposure to the air (Raffaelli & Hawkins 1999, Menge & Branch 2001). For example, in the summer on cold-temperate shores, daily maximum temperature can be 10 °C higher and algal desiccation during low tides four times higher at high elevations than at low elevations (Eckersley & Scrosati 2012). On NW Atlantic rocky shores, fucoid seaweed canopies *(Ascophyllum nodosum* and *Fucus* spp.) often cover the substrate extensively from low to high elevations in wave-sheltered habitats (Adey & Hayek 2005, Longtin et al. 2009; Fig. 1). Due to the limited aerial exposure at low elevations, fucoid canopies in such places have almost no influence on benthic temperature and do not affect the structure of benthic communities. However, with the longer aerial exposure at high and middle elevations, fucoid canopies limit the otherwise high thermal extremes and, in that way, increase the richness (number of species) of benthic communities (Watt & Scrosati 2013). As fucoid canopies do not affect benthic richness at low elevations, this paper evaluates the occurrence of a unimodal facilitation–stress relationship by testing the hypothesis that the positive effect of canopies on benthic richness is greater at middle elevations than at high elevations.

**Figure 1.**
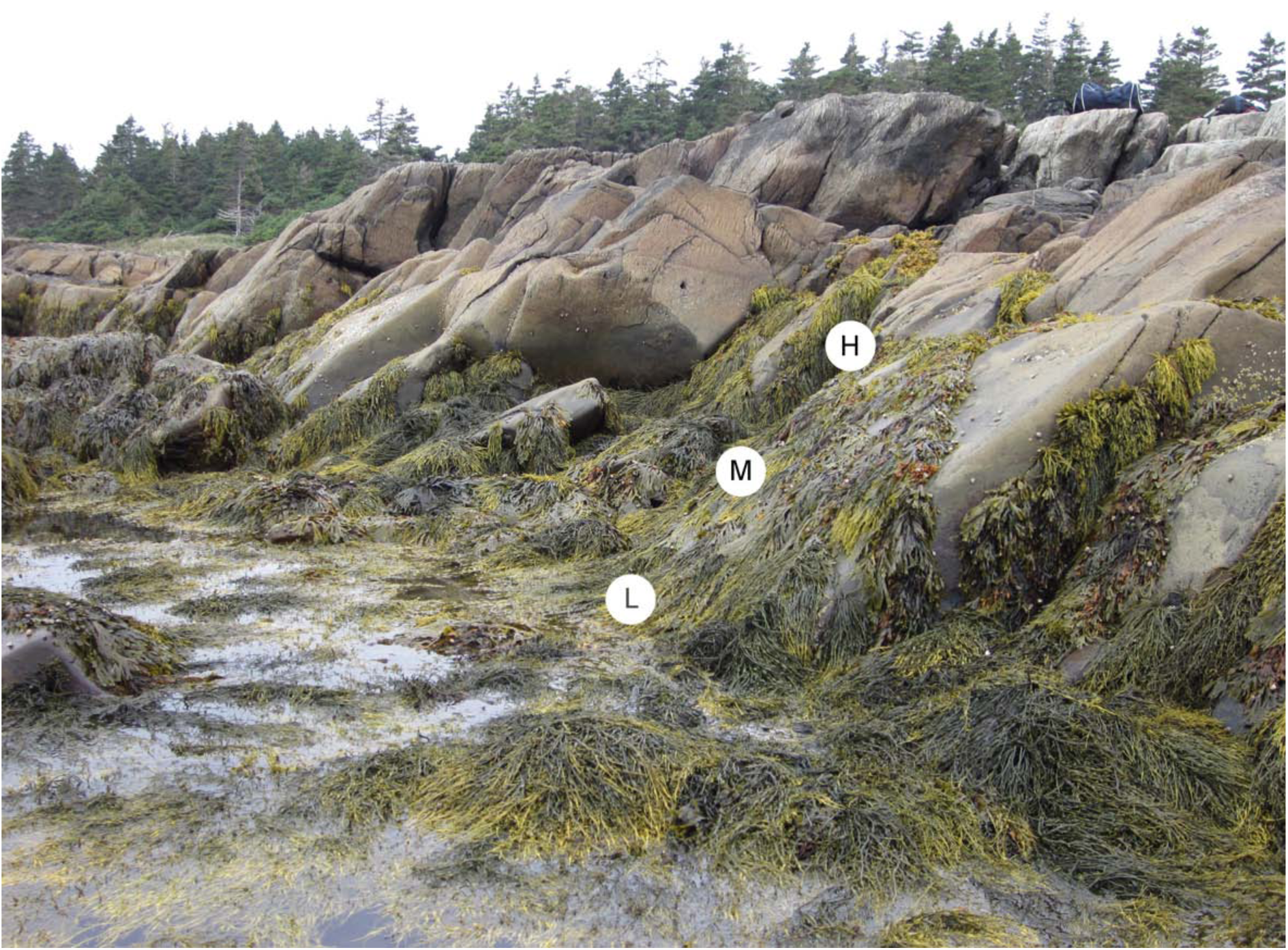
Wave-sheltered rocky intertidal habitat from the Atlantic coast of Nova Scotia. This view at low tide shows the extensive cover of fucoid algal canopies at high (H), middle (M), and low (L) elevations. Picture taken by Ricardo A. Scrosati.

## Materials and methods

This study uses the published data set (Watt & Scrosati 2014) that was produced by a mensurative experiment to show that fucoid canopies increase benthic richness at high and middle elevations while having no effects at low elevations (Watt & Scrosati 2013). The methodology of that study is described in the corresponding paper (Watt & Scrosati 2013), but it is summarized here to highlight the main aspects. The data were measured in wave-sheltered rocky intertidal habitats spanning 350 km of the Atlantic coast of Nova Scotia, Canada. These canopies have a similar composition of fucoid algae (Fucaceae) at low (0-0.5 m above chart datum), middle (0.5-1 m), and high (1-1.5 m) elevations, with a predominance of *Ascophyllum nodosum* (59-72 %) followed by *Fucus vesiculosus* (26-34 %) and three other species of *Fucus* (< 1-7 %). To assess whether canopy effects on benthic richness existed at each elevation zone, all algae and invertebrates found in replicate quadrats (20 cm x 20 cm) randomly placed at each zone were identified. For each elevation zone, canopy effects were looked for by comparing richness between two canopy cover treatments: low (0-40 %) and high (60-100 %) cover. Since fucoid canopies only affected richness at high and middle elevations (Watt & Scrosati 2013), only data for such elevations were necessary to test the hypothesis of the present study. The number of surveyed quadrats was 233 (low canopy cover) and 110 (high canopy cover) for high elevations and 111 (low canopy cover) and 234 (high canopy cover) for middle elevations. For these two elevation zones, a total of 16 seaweeds (excluding the fucoid species) and 41 invertebrates were identified (Watt & Scrosati 2013).

In this study, both the intensity and importance (Brooker et al. 2008) of whole-community facilitation by fucoid canopies are compared between high and middle elevations. Facilitation intensity was calculated for each elevation zone as the percent increase in richness between the low-cover and high-cover treatments. For high elevations, the 110 high-cover quadrats were randomly paired with 110 low-cover quadrats selected at random while, for low elevations, the 111 low-cover quadrats were randomly paired with 111 high-cover quadrats also selected at random. For each resulting pair of quadrats, the percent change in richness was calculated as {[(*S_H_ - S_L_*)/*S_L_*]*100}, where *S*_*H*_ was species richness in the high-cover quadrat and *S*_*L*_ was richness in the low-cover quadrat. Thus calculated, facilitation intensity was compared between high and middle elevations through a two-sample *t*-test (Howell 2002). The importance of facilitation was calculated for each elevation zone using the point-biserial correlation coefficient (*r*_**pb**_). To calculate *r*_**pb**_ for each zone, richness was considered as the dependent variable and the two canopy cover treatments were considered as the independent variable, coding low cover as “1” and high canopy cover as “2” (Fritz et al. 2012). The percent *r*_pb_^2^ was calculated to indicate the percentage of variation in richness that could be explained by canopy cover, which was considered as the importance of whole-community facilitation (*r*_**pb**_ being positive) relative to all other sources of variation in richness. The point-biserial correlation coefficient was compared between high and middle elevations using the *Z* test designed to compare two independent *r* values (Howell 2002). These analyses tested the hypothesis of this study at the patch (quadrat) scale. The difference in facilitation intensity between both elevation zones was also evaluated at the whole-habitat scale (Cavieres et al. 2016), for which the total number of species found only under high canopy cover, only under low cover, and in both cover treatments (Armas et al. 2011) was calculated for middle and high elevations using the information provided in Table 2 in Watt and Scrosati (2013).

## Results

At the patch scale, the intensity of community-level facilitation by fucoid canopies was, on average, 36 % higher at middle elevations than at high elevations, which was a significant difference (*t*_219_ = 2.25, *P* = 0.026; Fig. 2A). The importance of facilitation was also higher at middle elevations, as fucoid canopy cover explained 49 % (percent *r*_pb_^2^) of the observed variation in benthic richness at middle elevations and 38 % at high elevations. The point-biserial correlation coefficient (*r***pb**) was significantly higher at middle elevations than at high elevations (*Z* = 1.88, *P* = 0.030; Fig. 2B). At the whole-habitat scale, 26 of the 57 species (46 %) identified at middle elevations were only present under high canopy cover, while no species (0 %) were only present under low cover. At high elevations, however, just 11 of the identified 30 species (37 %) were only present under high canopy cover, while 3 of those 30 species (10 %) were only present under low cover.

**Figure 2.**
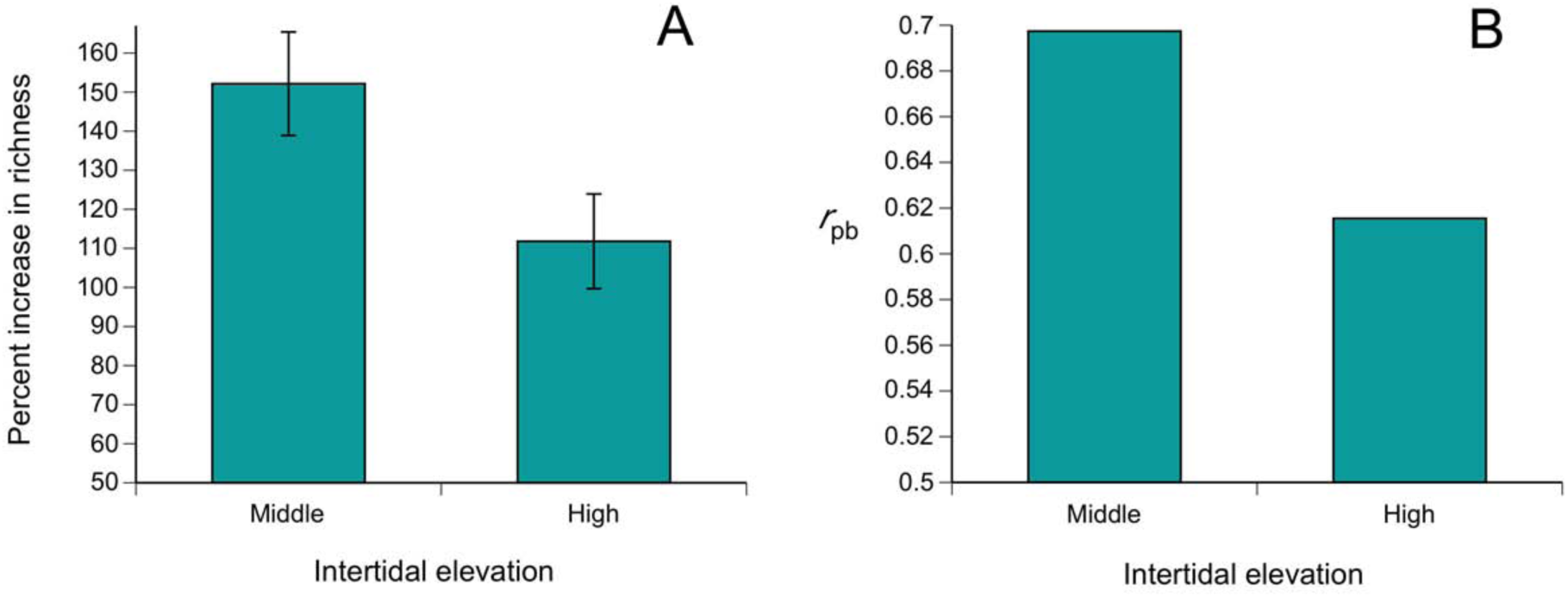
Community-level facilitation by fucoid algal canopies. (A) Percent increase in benthic species richness between quadrats with low and high canopy cover (facilitation intensity; mean ± SE) at middle and high intertidal elevations. (B) Point-biserial correlation coefficients (P < 0.001 in both cases) used to calculate the importance of facilitation at middle and high elevations. Both descriptors of facilitation differed significantly between middle and high elevations (see Results for details).

## Discussion

A recent study (Watt & Scrosati 2013) found that intertidal fucoid canopies increase benthic species richness at middle and high elevations. The present study has revealed that such effects are greater at middle elevations in terms of both intensity and importance. As fucoid canopies have no effects on benthic communities at low elevations (Watt & Scrosati 2013), these findings support the unimodal facilitation–stress hypothesis (Michalet et al. 2006, Holmgren & Scheffer 2010). In other words, community-level facilitation by intertidal fucoid canopies peaks at an intermediate level of environmental stress represented by middle elevations.

A variety of mechanisms have been proposed to explain the drop in facilitation intensity from intermediate to high stress levels (Michalet & Pugnaire 2016). When effects are analyzed at the community level (as in this study), a commonly proposed mechanism is the decreasing ability of facilitators to improve conditions strongly enough for some species towards the highest stress levels where the facilitators occur (Holmgren & Scheffer 2010, He & Bertness 2014, Michalet et al. 2014). For intertidal communities as a whole, physiological stress (mainly due to high temperature and desiccation during low tides) peaks at high elevations (Raffaelli & Hawkins 1999, Menge & Branch 2001). However, at high elevations, fucoid canopies were found to be unable to limit mean temperature as strongly as at middle elevations (Watt & Scrosati 2013). Thus, these observations lend support to the above explanation. Studies on the physiological influence of fucoid canopies on the benthic species found at high and middle elevations (currently lacking) could contribute to strengthen this view.

Another explanation for facilitation decreasing at high stress relates to structural changes in the facilitators. For example, Bonanomi et al. (2016) found a hump-shaped relationship between altitude on mountain sides (proxy for cold and wind stress) and the intensity of facilitation by cushion plants on associated plant richness.The decreasing facilitation at high altitudes seemed to result mainly from an increase in cushion compactness, which may have limited the ability of cushions to trap seeds of other plants and/or enable their root development (Bonanomi et al. 2016). This was not the case for intertidal fucoid canopies, however. These canopies are extensive but do not increase in compactness towards high elevations. Moreover, the canopies arise from relatively small holdfasts (the structures that keep algae attached to the substrate), leaving ample substrate for other benthic species to occur.

Another proposed explanation for the decrease in facilitation at high stress involves increasing competition. For example, when stress peaks due to intense water loss in the soil, canopy-forming plants may actually compete for water with the associated plants, which can limit or even eliminate facilitation (Holmgren et al. 2012, Michalet et al. 2014, Butterfield et al. 2016). However, this scenario is not applicable to the studied intertidal habitats, because benthic algae and sessile invertebrates are attached to solid bedrock in these places.

Regardless of the underlying mechanism, decreases in facilitation intensity from intermediate to high stress levels have been found for additional systems recently (de Bello et al. 2011, Koyama & Tsuyuzaki 2013, Castanho et al. 2015). Overall, these findings point to the more complex nature of the facilitation–stress relationship than originally envisioned. In this sense, the contribution of the present study is important because it is based on data for the entire community, including primary producers and consumers. This is relevant because most facilitation studies have investigated effects on a few associated species or, when looking at the multispecies level, often only on the assemblage of associated plants (Soliveres et al. 2015, Bonanomi et al. 2016, Cavieres et al. 2016, López et al. 2016). Recent studies are recognizing the need to evaluate facilitation effects at the whole-community level, including plants and animals, to develop a broader conceptual understanding of the facilitation–stress relationship (Ruttan et al. 2016, Lortie et al. 2016).

